# Novel carbon nanoparticles (CNPs) from *Catharanthus roseus* petals for metal ion sensing and root growth modulation

**DOI:** 10.1101/2024.03.18.585564

**Authors:** Satya Omprabha, Charli Kaushal, Nilesh D. Gawande, Raviraj Barot, Dhiraj Bhatia, Subramanian Sankaranarayanan

## Abstract

Nanoparticles are of major interest nowadays due to their distinctive properties. In this study, we present a microwave-assisted green synthesis of carbon nanoparticles from the *Catharanthus roseus* petals. The nanoparticles exhibited a fluorescence emission in the 300 to 400 nm wavelength region and emitted a characteristic blue fluorescence. The extensive characterization using Bio-AFM showed that the size of nanoparticles was 25.38 nm. M-XRD and Zeta potential analysis confirmed that the nanoparticles were amorphous and exhibited a negative surface charge. The surface functional groups on the nanoparticles were highlighted by FTIR analysis. Metal sensing affinity of the synthesized CNPs and their effect on the plant growth was tested. These CNPs demonstrated the effective sequestration of metallic ions such as copper (Cu^2+^), iron (Fe^2+^), and nickel (Ni^2+^), indicating their potential to be used as biosensors in metal detection. It was found that the effect was proportionate to the *Catharanthus roseus* petal nanoparticles (CRPNP) concentration, where it exhibits a great affinity for Fe^2+.^ The effect of the nanoparticles analyzed on the root growth of *Arabidopsis thaliana* showed that CRPNP suppresses the primary root and lateral root numbers. These results demonstrate the use of *C. roseus* synthesized CNPs in sustainable agriculture by targeting nutrient delivery.

## Introduction

The concept of “nanotechnology” was first introduced by physicist Richard Feynman in his famous 1959 lecture titled “There’s Plenty of Room at the Bottom.”(I. Khan et al., 2019). Nanomaterials have a size or one of the dimensions ranging from 1 to 100 nm (Baig et al., 2021). Due to their distinctive optical, electronic, magnetic, mechanical, and chemical properties, which are primarily derived from the quantum confinement effect and large surface-to-volume ratios, nanostructured materials with controllable size, shape, crystallinity, and tunable surface functionalities have attracted extensive research attention (Li et al., 2015). Recent advances in this field have introduced various sources for the preparation of nanoparticles (NPs) with diverse applications (Sharma et al., 2017). Green synthesis is one such method that offers numerous advantages, including lower toxicity, cost-effectiveness, and scalability. Moreover, plants contain many phytochemical compounds such as phenolics, terpenoids, polysaccharides, and flavonoids that possess oxidation–reduction capabilities, which are preferably utilized for the green synthesis of nanoparticles (Vijayaram et al., 2023). The biological molecules present in the phytochemicals, used as both reducing and capping agents provide an advantage in preventing the agglomeration of NPs, reducing toxicity, and improving antimicrobial action (Thapa & Roy Choudhury, 2021). Numerous sources may act as precursors for green synthesis such as fruits, fruit juices and peels (Lu et al., 2012), beverages (Zhu et al., 2012), animal and animal derivatives (Wang et al., 2012), flour and bakery products (Sk et al., 2012), human derivatives (D. Sun et al., 2013), and vegetables, and spices (Lu et al., 2013).

Recent developments in this field have enhanced the applications of nanomaterials in tissue engineering, cancer therapy, protein detection, drug delivery, etc (Salata, 2004). Their unusual interactions with biomolecules both outside and inside of cells, due to their unique features, have significantly improved cancer detection and therapy (Cai, 2008). Several clinically approved nanomedicines are used to treat diseases such as breast cancer, ovarian cancer, Kaposi’s sarcoma, etc (Blasiak et al., 2013). Biosensing is also one of the major applications in nanotechnology, and it includes colorimetric sensing, fluorescence sensing, electrochemical sensing, and other sensing systems (De et al., 2008).

Nanoparticles are a major choice for metal detection nowadays. Various studies have explored the properties of metal sensing in nanoparticles (Cecchini et al., 2014; Priyadarshini & Pradhan, 2017; Uddandarao & Balakrishnan, 2017; H. Yang et al., 2013). For instance, carbon dots of 2.6 nm size synthesized by one-step microwave hydrothermal method from China grass carp scales consisting of a large number of cysteines containing sulfhydryl groups were responsible for the detection of Hg^+2^ demonstrating it as a usable sensitive probe to detect Hg^+2^ (Liu et al., 2019). The pomelo peel was used to create fluorescent CNPs with a detection limit of 0.23 nM for Hg^+2^ (Lu et al., 2012). Other heavy metals, such as Cd+2 ions, were detected by AgNPs derived from neem seeds (Pawar et al., 2019). Metals ions Pb^+2^, Cu^+2^, and Ni^+2^ were also detected by ZnS Gold NPs formed from the fungi *Aspergillus flavus* (Uddandarao et al., 2019a). High-quality bismuth nanoparticles have been used to prepare electrodes and utilized to detect Cd^+2^ and Pb^+2^ (H. Yang et al., 2013). Various amino acids have been used to synthesize carbon quantum dots (CQDs) to detect heavy metal ions, out of which CQD prepared from Leucine showed selective detection for Fe^+3^ ions (Šafranko et al., 2023).

Several studies have been conducted to determine the effect of nanoparticles on the modulation of plant growth. For instance, CuNPs of 50 nm size were found to reduce the *Cucurbita pepo*’s root length growth by 77% and biomass by 90% (Stampoulis et al., 2009). The effect of silver NPs synthesized from *Acalypha indica* Linn. leaves extract studied on the root, stem, and leaf tissues of *Bacopa monnieri* (Linn.) showed an insignificant reduction in the shoot and root lengths, affecting the air chambers in the root cortex, and xylem elements in the stem (Krishnaraj et al., 2012). Metal NPs have been shown to increase the length of the above-ground part of the seedlings of *Triticum aestivum L*. and *Hordeum vulgare L*. by 8-22%, with the alleviation in the chlorophylls and carotenoids levels (Hoang et al., 2022). NPs synthesized from SiO_2_ at different concentrations have affected plant growth in many ways, promoting seed germination in tomatoes, seed sprouting in Maize, and enhanced antioxidant activity in tomatoes and squash. Similarly, ZnO NPs have a stimulatory effect on the seed germination of cucumber, alfalfa, and tomatoes while enhancing plant development, biomass, leaf pigments, and protein content in Cluster beans (M. R. Khan et al., 2019). Various studies have determined that nanoparticles may have beneficial and detrimental effects on plants, depending on their size, concentration, chemical makeup, zeta potential, stability, and shape (Rastogi et al., 2017).

*Catharanthus roseus*, also known as *Vinca rosea*, is a dicotyledonous angiosperm belonging to the Apocynaceae family. It is native to the Indian Ocean Island of Madagascar but can be found in many tropical and subtropical regions worldwide (Verma & Mishra, 2017). *C. roseus* is a traditional remedy for wasp stings, central nervous system (CNS) suppression, and muscular discomfort. Besides treating sore throats and mouth ulcers, it has also been used to cure nose bleeds and gum hemorrhage. (Rajeswari, 2013). Silver nanoparticles prepared from a mixture of aqueous silver nitrate and flower extract of *C. roseus* exhibit a potential antibacterial activity against *E. coli, P. putida, S. aureus, K. pneumonia*, and *B. subtilis* (Alwala et al., 2014). Silver nanoparticles derived from *C. roseus* flowers are hemolytic in a concentration-dependent manner (Raja et al., 1970). Iron nanoparticles synthesized from the leaves of *C. roseus* also displayed antibacterial activity against *E. coli* and had a 50% dye-degrading ability against Methyl orange (Roy et al., 2022). These studies showcase the potential of *C. roseus* nanoparticles in various applications.

In the present study, novel nanoparticles have been synthesized through a single-step microwave-assisted reaction using a petal extract of *Catharanthus roseus* (CRPNP). An extensive characterization of CRPNPs was performed using various spectrophotometric and spectrofluorometric methods. CNPs were utilized to evaluate the metal-sensing properties of iron (Fe^2+^), nickel (Ni^2+^), and copper (Cu^2+^). The effect of the CRPNPs was also analyzed on the root growth of *Arabidopsis thaliana*. Our study demonstrated that CRPNP possesses the highest sensing ability for Fe and has a detrimental effect on the root growth of *Arabidopsis thaliana*.

## Materials and methods

### Synthesis of petal-derived nanoparticles

Fresh flowers of *Catharanthus roseus* species were collected from the Indian Institute of Technology Gandhinagar (IITGN) campus, Palaj, Gujarat, India. A quantity of 5 grams of cleaned petals was crushed using 30 mL of ethanol, followed by centrifugation at 15,000 rpm and 25°C for 10 minutes. The supernatant was filtered through a 0.22 μ membrane filter (Tarsons®) and stored at room temperature for further experimental purposes. The petal extract of a volume of 5 mL was mixed with 10 mL of ethanol and microwaved for 8 minutes until the liquid phase evaporated completely. NPs obtained were then dissolved into 10 mL of ethanol after sonication for 2 minutes. The solution was again filtered using a 0.22 μm membrane filter and stored at room temperature. These NPs were called CRPNPs, representing the Catharanthus roseus petal-derived nanoparticles.

### Metal detection assay

Working solutions of CRPNPs were prepared at concentrations of 50, 100, and 150 μg/mL from the stock solution of 10 mg/mL and stored at room temperature. The metal ions of Iron (Fe^2+^), Nickel (Ni^2+^), and Copper (Cu^2+^) were prepared using their salts at a concentration of 100 μM/mol dm^-3^. Mixtures of metal ions (100 μL) and NPs (900 μL) were vortexed for 30 seconds and were incubated at room temperature for 40 minutes. The emission spectra of CRPNP-metal ion mixtures were recorded at an excitation wavelength of 400 nm using a UV-Vis spectrofluorometer after the incubation at room temperature. The blank samples for each of the concentrations of CRPNP tested had the following values of fluorescence intensity: 50 μg/mL-463.717 nm, 100 μg/mL-792.069 nm, and 150 μg/mL-1061.92 nm, which were considered as a reference to calculate the fluorescence response. Fluorescence spectra of CRPNP with each metal ion were analyzed in Origin 2017 Software, and fluorescence response was quantified using the following formula:

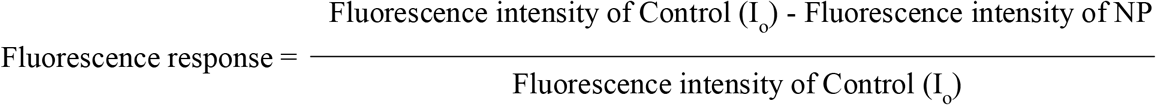

### Plant growth assay

Seeds of *Arabidopsis thaliana* wild-type genotype *Col-0* were sterilized with 70% ethanol for 5 minutes, followed by a 5-minute wash with 4% bleach. Seeds were washed 4-5 times using sterilized Milli-Q water and were germinated on the sterile half-strength Murashige Skoog (MS) agar media plates containing 0.22% MS salt, 0.05% MES monohydrate, 2% sucrose, and 0.8% plant agar (pH 5.7 adjusted with 1M KOH). Plates were incubated at 4°C for two days before being moved to the plant growth chamber under long-day conditions and a light intensity of 130 mol/m^2^/s. Three days after germination, the seedlings were transferred to the control half MS plates (without CRPNP) and treatment plates, half MS plates supplemented with 5 mg/mL of CRPNP. The effect of the CRPNP on the root growth was analyzed on day seven after the treatment.

### Characterization of CRPNP

Various methods were employed for the extensive characterization of the synthesized CRPNPs. The functional groups on the surface of NPs were identified using a Fourier-transform infrared spectrometer (FTIR, Perkin Elmer^®^ Spectrum Two FTIR Spectrometer). The size of NPs was determined using a Bio-Atomic Force Microscope (Bio-AFM, Bruker^®^ Nano wizard Sense AFM). The nature of NPs was determined using multipurpose X-ray diffraction (M-XRD, Rigaku^®^ SmartLab 9KW). The optical properties of the NPs were analyzed using a UV-VIS spectrophotometer (Analytika Jena SPECORD^®^ 210 PLUS) and a spectrofluorometer (Jasco^®^ Spectrofluorometer FP-8300). The surface charge on the CRPNPs was determined using DLS/ZETA Sizer (Nano ZS, Malvern^®^ Instruments).

### Statistical analysis

Student’s t-test was used to determine the significant differences between the control and CRPNP treatments. The graphs were plotted using Prism version 8.0.2.

## Results

We have synthesized the nanoparticles (NPs) from the petal extract of *Catharanthus roseus*, characterized the CRPNPs, and studied the application of these particles for metal biosensing ability and their effect on the root growth of *A. thaliana*. The outline for the synthesis of nanoparticles and their application is illustrated in **Fig. 1** and **Fig. 2**.

**Fig. 1.**
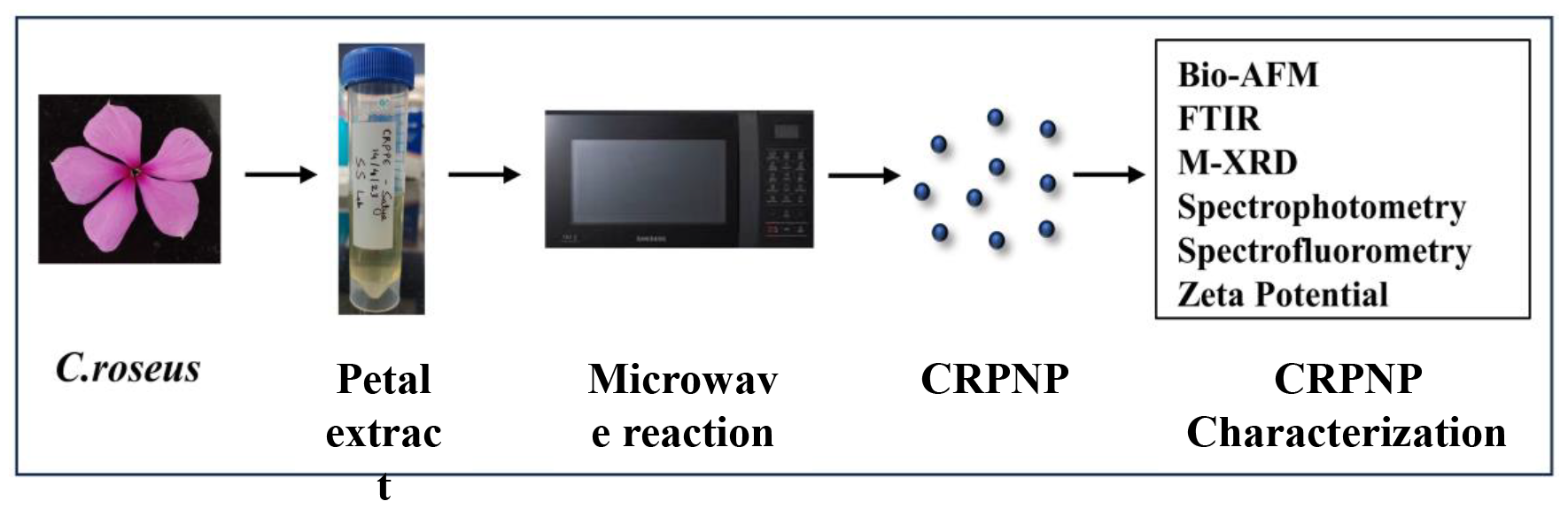
Synthesis of petal-extract derived nanoparticles from *Catharanthus roseus* and its characterization. CRPNPs were synthesized from the petals of *C*.*roseus* using the microwave method and further characterized by various characterization techniques.

**Fig. 2.**
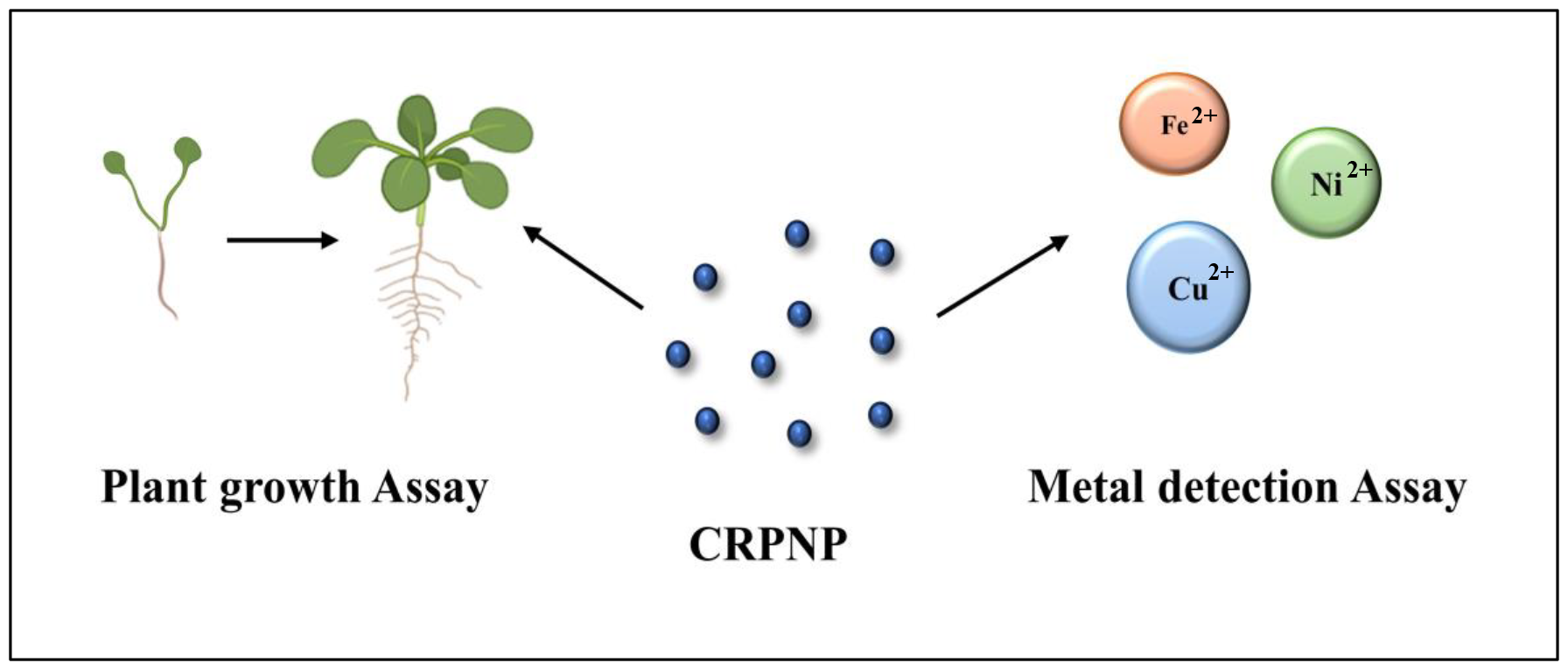
Application of CRPNPs in this study. The CRPNPs were tested for their biosensing effect on metals, iron (Fe^2+^), copper (Cu^2+^), and nickel (Ni^2+^). Also, the changes in root growth of *A. thaliana* were recorded to determine the impact of nanoparticles.

### NPs synthesized from *Catharanthus roseus* petals have a size of less than 50 nm

The size profile of CRPNPs derived from petal extract of *C. roseus* analyzed by Bio-AFM imaging technique revealed that these were two-dimensional NPs with an average diameter of 25.38 ± 1.44 **(Fig. 3A)** and height of 2.98 ± 0.15 **(Fig. 3B)**.

**Fig. 3.**
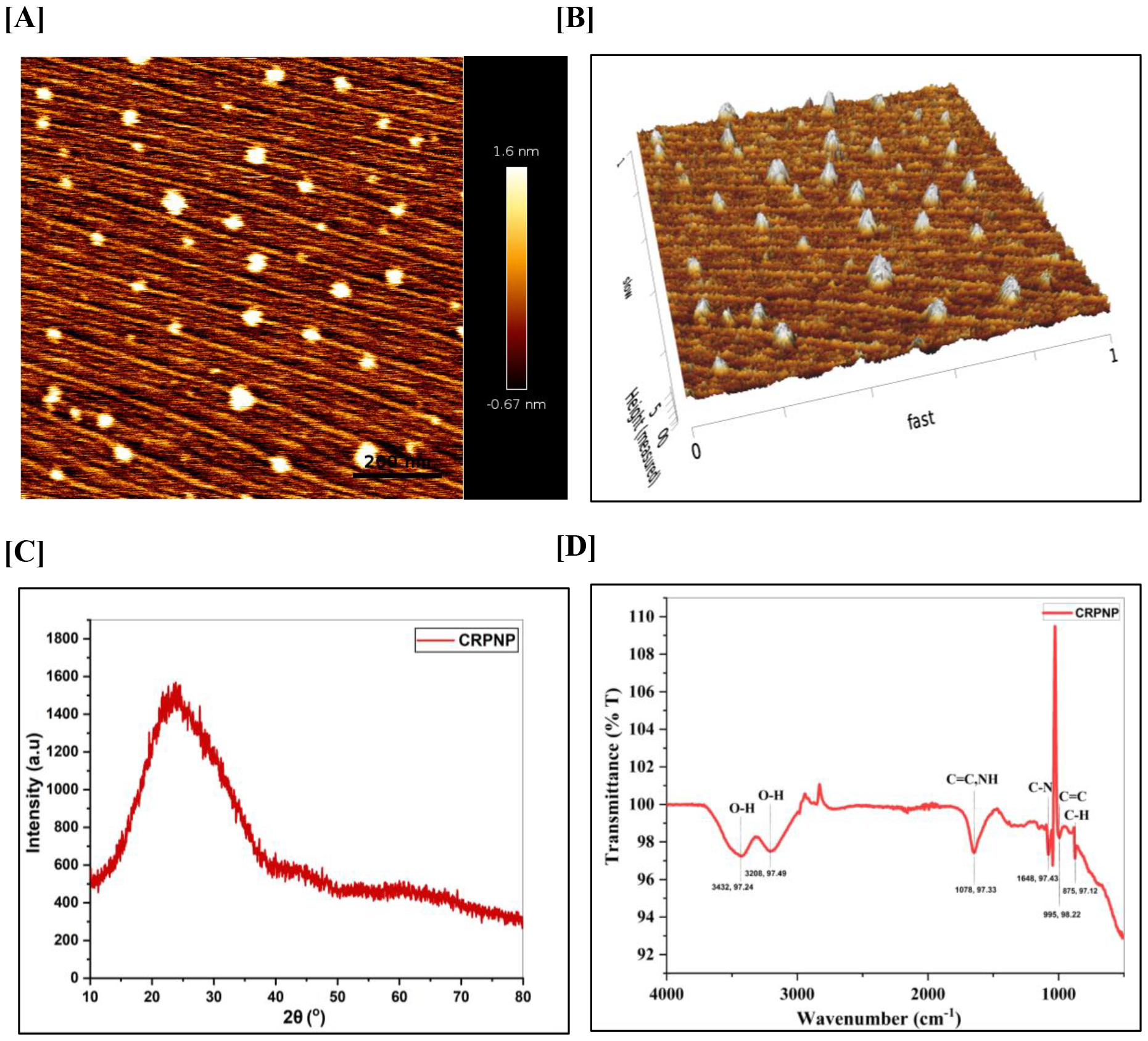
Structural characterization of synthesized nanoparticle from the petal extract of *Catharanthus roseus*. The size and height of CRPNP were analyzed using Bruker^®^ Nano Wizard sense AFM, with the scale bar 200 nm for **(A)** and 5nm for **(B)** respectively. **(C)** M-XRD analysis was carried out using igaku^®^ SmartLab 9KW, with a scan rate of 4° per minute and a range of 10° to 80° of 2θ X-ray diffraction angle. **(D)** FTIR graph shows functional groups present over CRPNP, analyzed using Perkin Elmer^®^ Spectrum Two FTIR Spectrometer. Graphs were plotted by using the Origin 2017 version.

The M-XRD (Multipurpose X-ray diffraction) profile of CRPNPs exhibited a broad peak centered around 23.84° (2θ). The *d*-spacing value (Å) remained at 3.72944 **(Fig. 3C)**.

The absence of any significant strong peaks suggested CRPNPs were amorphous. Also, the zeta potential analysis of petal-derived NPs showed that they were negatively charged at – 3.92 mV. The presence of different functional groups belonging to certain classes was determined through an FTIR analysis suggesting broad peaks at 3432 cm^-1^, corresponding to O-H stretching vibrations belonging to the class of alcohol and at 3208 cm^-1^ belonging to carboxylic acid or alcohol. The stretching vibrations of C=C and bending vibrations of N=H groups were observed at 1648 cm^-1^, the former representing the class of alkene and the latter of amine. CRPNP also exhibited a peak at 995 cm^-1^, suggesting the presence of bending vibrations of functional group C=C, but it belonged to the class of allenes, also classified as cumulated dienes. The stretching vibrations of C-N were observed at a medium peak at 1078 cm^-1^, which corresponds to the class of amine. Also, a typical strong peak at 875 cm^-1^ represented the bending vibrations of C-H belonging to 1,2,4-trisubstituted or 1,3-disubstituted classes **(Fig. 3D)**.

### CRPNPs emitted blue fluorescence under UV

The UV-visible absorption spectrum of petal extract CRPNPs displayed two peaks, one with the intensity of 0.7 a.u between 250 nm and 300 nm in the mid-energy region **(Fig. 4A)**. This peak corresponds to the surface states with the presence of C=C structures and the π-π* transitions of C=C bonds. The other one was a broad peak with an intensity of 0.5 between the wavelength 300 nm and 400 nm in a low-energy region. This peak represents the surface states with C=C-C=C (allenes) structures and π_2_-π_3_* transitions. The photoluminescence spectra of synthesized petal extract CRPNPs demonstrated a wide-band emission of 250 nm to 600 nm **(Fig. 4B)**. Under natural light or daylight conditions, the synthesized CRPNPs displayed a light-yellow color in an aqueous solution **(Fig. 4C)**, whereas emitted blue color fluorescence under UV excitation **(Fig. 4D)**. The highest fluorescence emission intensity observed was at 417 nm of 6544 a.u when photoexcited at a wavelength of 300 nm.

**Fig. 4.**
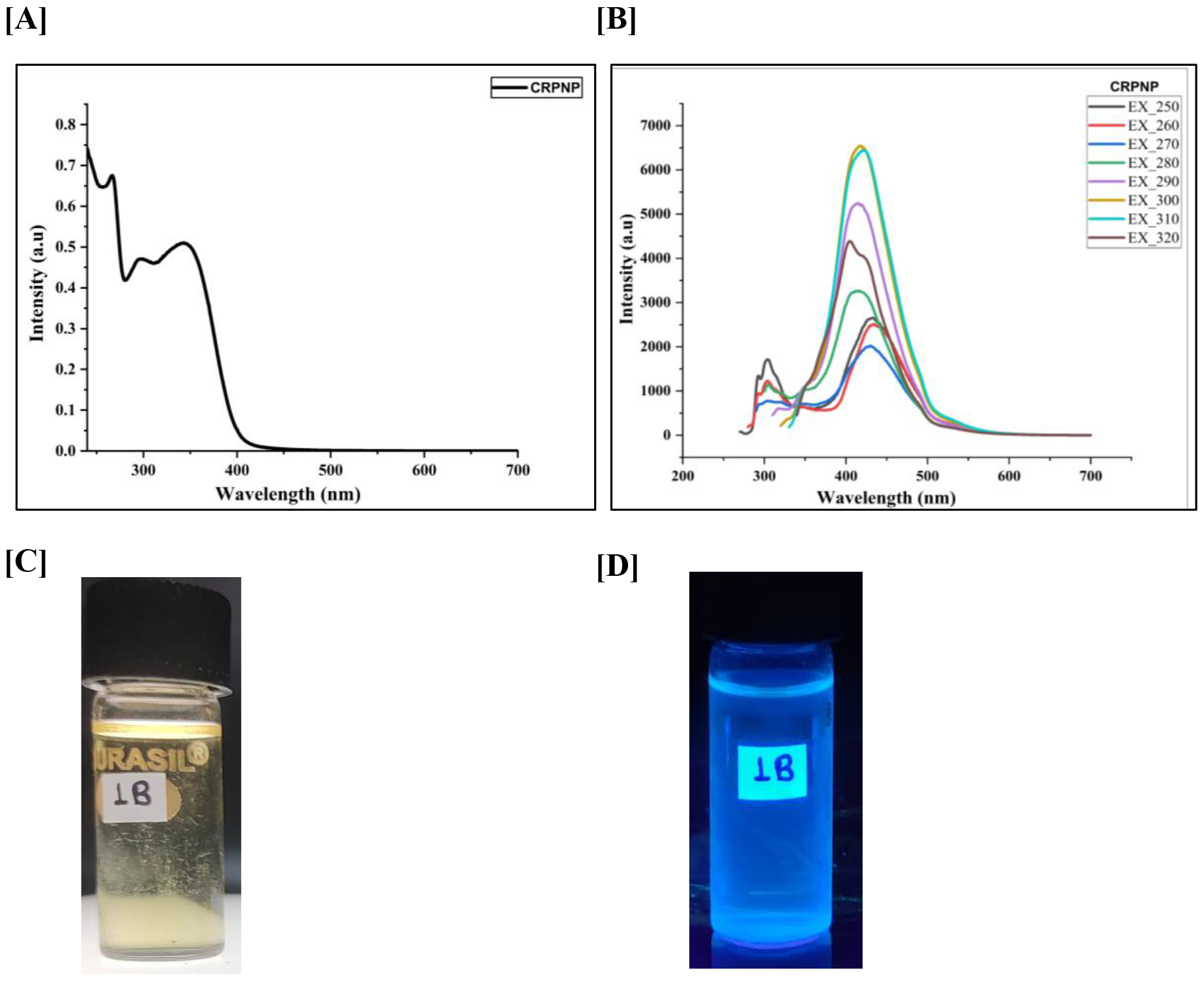
Characterization of optical properties of CRPNP. **(A)** Spectrophotometric analysis and spectrofluorometric analysis. **(B)** of petal-derived NP which is labelled as 1B was performed using Analytika Jena SPECORD^®^ 210 PLUS spectrophotometer and JASCO^®^ spectrofluorometer FP-8300 respectively. **(C)** CRPNP was visualized in white light and under UV light **(D)**. The graphs were plotted using Origin Software 2017 version.

### CRPNPs showed a significant sensing effect by sequestration of metallic ions

The fluorescence response recorded for all three concentrations of CRPNPs with metal ions was in the order of Ni^2+^ < Cu^2+^ < Fe^2+^, where an increase in the concentration of CRPNPs increased the fluorescence response **(Fig. 5 and Supplementary Table S1)**. The fluorescence of the NPs with the metal ions was of lower intensity than the control (without NP) when CRPNPs were excited at 400 nm. CRPNPs showed higher affinity at the concentration of 150 μg/mL for each metal. The difference in fluorescence response at the concentration from 50 to 100 μg/mL was surprisingly decreased and significantly increased from 100 to 150 μg/mL. The CRPNP’s behaviour towards the metal ions suggests it has a good biosensing ability and sensitivity for heavy metals at higher concentrations.

**Fig. 5.**
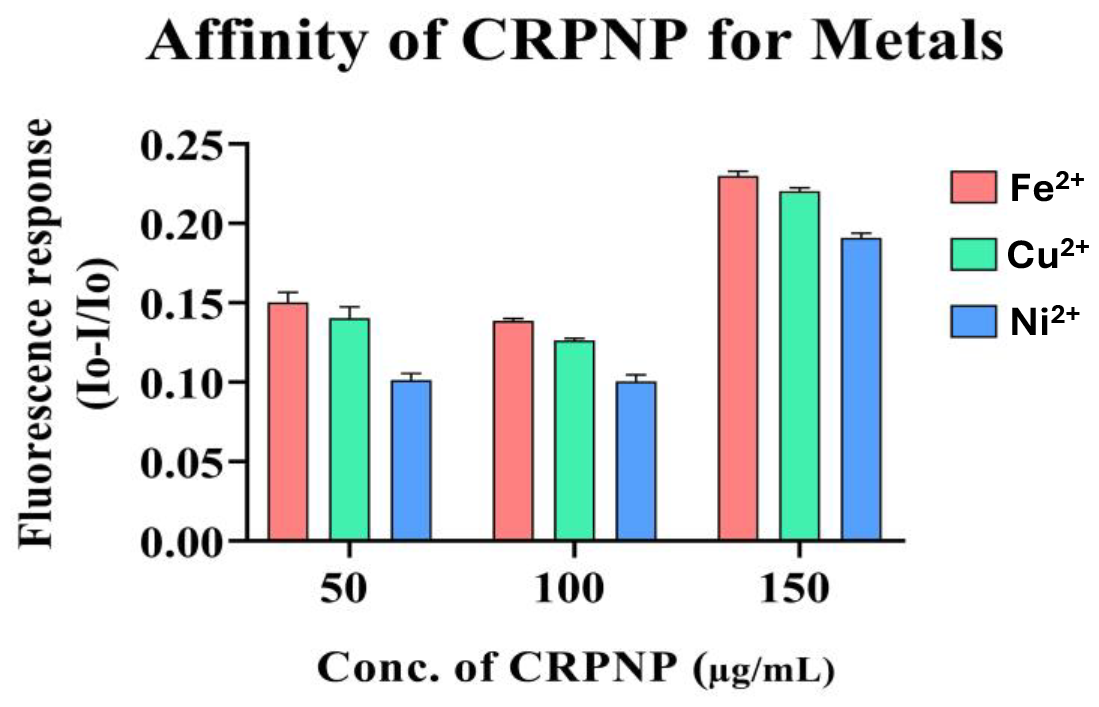
Determination of affinity of CRPNP for metals. The affinity of the CRPNP for metals such as Iron (Fe^2+^), Copper (Cu^2+^), and Nickel ((Ni^2+^) was determined at different concentrations of CRPNP such as 50 μg/mL, 100 μg/mL, and 150 μg/mL. Fluorescence response is increased in a concentration-dependent manner. The response was recorded using Jasco^®^ spectrofluorometer FP-8300. Graphs were plotted using Prism Version 8.0.2.

### CRPNPs suppressed the primary root growth and lateral root numbers

The root growth analyzed seven days after treatment with 5 mg/mL of CRPNP in *A. thaliana* displayed significant suppression of the primary root lengths and lateral root numbers **(Fig. 6A-D and Supplementary Table S2A-B)**. In comparison to the control, the primary root growth had enhanced suppression by 7.30%, and the number of lateral roots was suppressed by 25.38% (**Fig. 6C** and **D**). These results suggested that CRPNP has a very low effect on the suppression of primary root growth at a concentration of 5 mg/mL while it suppresses the lateral root numbers to a greater extent.

**Fig. 6.**
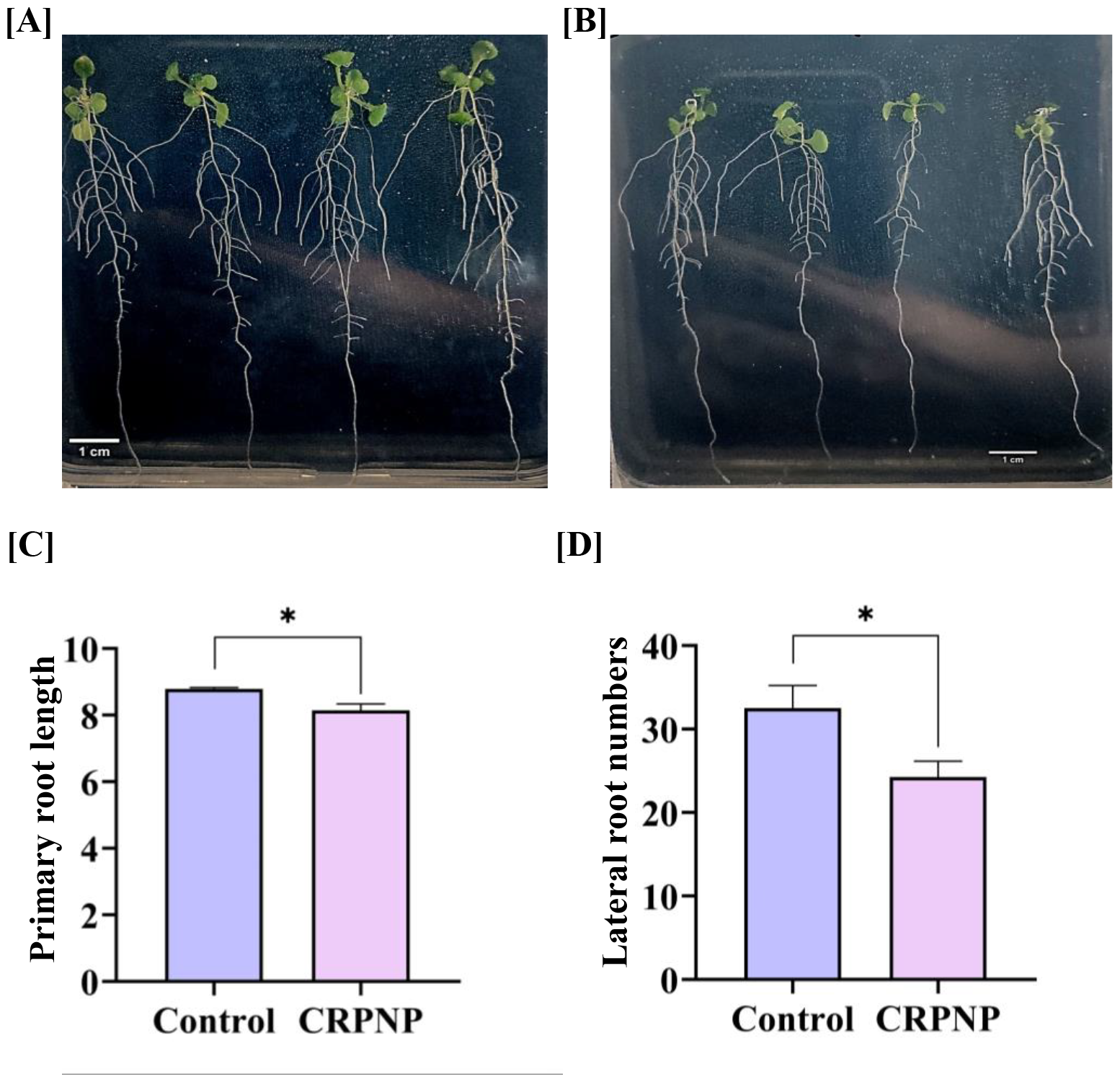
Effect of CRPNPs treatment on primary and lateral root growth of *A. thaliana*. **(A)** Control treatment (without CRPNP) and **(B)** 5 mg/mL *C. roseus* petals derived CNP treatment (CRPNP). **(C)** Suppression of primary root (PR) length, and **(D)** Suppression of lateral root (LR) number in response to CRPNP treatment. The effect of the CRPNP treatment on the PR and LR root growth is determined after day seven of the plant treatment. Significant differences between the treatments were analyzed using Student’s t-test. Graphs were plotted using Prism Version 8.0.2. Asterix (*) indicates that the significance level of <0.05.

## Discussion

### Morphological and structural specifications vary with chemical composition

*C. roseus* has been known for its antioxidant, anthelminthic, anti-hyperglycaemic, in vivo antidiarrheal, antimicrobial, antineoplastic, and antidiabetic effects (Rajeswari, 2013). It consists of various chemical compounds such as alkaloids-vinblastine, and vincristine is mostly found in the aerial parts of the plant and widely used for neoplasm treatment; Ajmalicine, found in the roots, is beneficial in treating circulatory illness (Aslam et al., 2010). We have reported the synthesis of NPs from the petal extract of *Catharanthus roseus*, CRPNP. Plant sources have been utilized as capping and reducing agents in the preparation of NPs with metals as doping agents. Silver nanoparticles (AgNPs) from the leaf extracts of *P. frutescens* (Tavan et al., 2023), flower extracts of *Catharanthus roseus* (Kandiah & Chandrasekaran, 2021), and gold nanoparticles (AuNPs) from the leaf extract of *Ceiba pentandra* (Aji et al., 2022), and many more have been prepared. Whereas crude extracts from various plant products have been used to prepare NPs without any doping agents, such as from cabbage juice (Alam et al., 2015), orange juice (Sahu et al., 2012), lemon juice (Ding et al., 2017), peel of *Trapa bispinosa* (Mewada et al., 2013), etc. Here, in our study, we demonstrate a similar method of synthesizing NPs from petal extract of *C. roseus* without doping with additional substrate. The nature of the NPs depends on the *d* spacing value, as it increases the amorphous nature, indicating more oxygen-containing groups (Sahu et al., 2012). The presence of O-H groups in CRPNPs, as confirmed by FTIR analysis, also ensures that the M-XRD analysis results in an amorphous nature. The characteristics and sizes of NPs depend on the techniques used for forming them. Two major synthesis techniques have been utilized to date for the formation of NPs from plant extracts, which are hydrothermal and microwave-assisted methods (Abid et al., 2022). Microwave-assisted synthesis is generally faster, cleaner, and more economical than traditional hydrothermal methods (G. Yang & Park, 2019). The size of NPs synthesized using the microwave-assisted method depends on the reaction time, which can be as small as 2 nm within 10 minutes of heating (Gerbec et al., 2005). After heating the petal extract of *C. roseus* for 8 minutes, we obtained the size of CRPNP to be 25.38 nm on average. The smaller size of NPs is usually beneficial for many application purposes.

### CRPNPs show concentration-based effects in metal biosensing

The ability of NPs to function as a biosensing element for metals has been investigated widely. Because of its high selectivity and sensitivity, the utilization of biogenic nanomaterials produced using straightforward techniques, such as green synthesis, for the detection of harmful heavy metal contamination is a viable strategy (Nayak et al., 2022). As a result of their ease of bioaccumulation and biomagnification in the food chain, heavy metals have been found to harm the health of all life forms (Uddandarao et al., 2019b). So, we studied the fluorescence-based metal detection technique using synthesized CRPNP for heavy metals Fe^2+^, Cu^2+^, and Ni^2+^. The affinity of CRPNP for metal sensing may be due to the stretching and bending vibrations of functional group elements present on its surface belonging to p-block and s-block (C, N, O, and H) as analyzed by FT-IR. CRPNP efficiently interacts with the metals Fe^2+^, Cu^2+^, and Ni^2+^ in decreasing order as d-block elements are transitional and can attain properties of both p and s blocks (Albrecht, 2010). The effect was found to be dependent on the concentration of CRPNP, where it shows the highest affinity for Fe^2+^, having a fluorescence response of 15%, 13.8%, and 22.9% at concentrations of 50 μg/mL, 100 μg/mL, and 150 μg/mL. Similarly, the response displayed for Cu^2+^ at increasing concentrations was 14%, 12.6%, and 22%, and those for Cu^2+^ were 10.1%, 10%, and 19% **(Table 4)**. An average difference of 2% has been observed for the transition from 50 μg/mL to 100 μg/mL, whereas a significant increase in the fluorescence response has been observed in the transition from 100 μg/mL to 150 μg/mL.

### CRPNPs exhibit a detrimental effect on primary and lateral roots

Depending on the characteristics of the nanomaterials, the method of administration, and the type of plant, researchers concluded that nanoparticles could have either good or detrimental effects on plant growth and development (Nair, 2016). The maximum degree of growth promotion and lowest levels of oxidative stress were seen following treatment with the decahedral AgNPs (45 ± 5 nm). In contrast, the highest degree of cotyledon growth inhibition and highest levels of oxidative stress were seen following treatment with the spherical AgNPs (8 ± 2 nm) (Syu et al., 2014). The tiny size of the nanoparticles enables them to penetrate through biological membranes, collect in the interior environment, integrate with proteins, and subsequently alter biological activities (Hoang et al., 2022). Therefore, this evidence suggests that the size of NPs plays a significant role in the alteration of plant’s physiological or morphological properties. In this study, the petal extract-derived CRPNPs having the size of 25.38 ± 1.44 nm exhibited a detrimental effect on the root growth where primary root length and lateral root numbers were suppressed. This suggests that CRPNP plays a critical role in modulating the phenotypic characteristics of the plants, such as root growth.

### Molecular and hormonal signaling may be the key to the inhibitory effect of CRPNP

A cascade of events begins when NPs enter the plants – NPs penetrate the cell wall, get internalized by the plants, interact with different processes inside the cell, and result in 2 types of events: (i) provide micronutrients or through any other mechanism supports the plant growth, and (ii) interacts with the changing phytohormone levels, oxidative burst, oxidation of proteins, lipids, and nucleic acids, alteration in plant metabolism, and acts by treating them with changing the plant’s morphology where the plant survives but with a decrease in plant growth (Goswami et al., 2019). This suggests that NPs interfere in molecular and hormonal signaling during their treatment of plants. Many possibilities may occur for the phenotypic expression of the plant after treatment with CRPNPs. We hypothesize that the inhibition in the root growth after the CRPNP treatment is due to the interruption in the auxin signaling pathways, which causes the decrease in the auxin levels in the roots and hampers the root growth of the plants. Similar responses have been studied in *A. thaliana*, where the polyvinylpyrrolidone-coated silver nanoparticles (PVP-AgNPs) suppressed the root gravitropism and downregulated the auxin receptor-related genes in NPs treated plants (Sun et al., 2017). However, the nanoparticles synthesized from ZnO (ZnONPs) have been shown to affect multiple hormones (Vankoba et al., 2017). However, this hypothesis further requires experimental validations such as gene expression analysis for hormone-signaling-related genes such as auxin.

## Conclusion

In conclusion, we have synthesized petal-extract-derived NPs from *Catharanthus roseus* using a microwave-assisted technique. The UV-Vis spectrophotometric and spectrofluorometric analyses confirmed that CRPNP displays blue emission when subjected to UV illumination. CRPNPs were amorphous and negatively charged. It showed metal biosensing ability when tested against three heavy metals: Fe^2+^, Cu2^+^, and Ni2^+^. CRPNPs had an inhibitory effect on the primary root length and lateral root number when analyzed using *A. thaliana*. In the future, investigations could be conducted to explore additional applications for CRPNPs, such as testing metal contamination in water and minimizing metal toxicity in plants by supplementation into the soil.

## Acknowledgment

We thank the Indian Institute of Technology Gandhinagar for the internship opportunity for SO. We thank GSBTM for a post-doctoral fellowship to CK and IITGN for a post-doctoral fellowship to NG. This work was supported by a DBT Ramalingaswami Re-entry fellowship grant and a start-up grant from the Indian Institute of Technology Gandhinagar to SS.

## Author Contributions

SS conceived and designed the research. SO synthesized and characterized the CNPs. SO and CK performed experiments on the effect of CNPs on root growth in Arabidopsis. NG and RB assisted in the characterization, and SS supervised the experiments. SO, CK, and NG analyzed the data. SO wrote the manuscript, and NG, CK, and SS proofread and edited it. DB provided technical suggestions for experiments. All authors read, discussed, and approved the manuscript.

